# Temperature-induced shift in the rare microbiota of the sponge *Haliclona*

**DOI:** 10.64898/2026.04.22.720130

**Authors:** Tyler J. Carrier, Frank Melzner, Sabrina Jung, Ute Hentschel

## Abstract

Ocean warming is currently leading to distributional shifts of species and an alternation of coastal communities. Vulnerable species that are most sensitive to ocean warming are able to use several acclimation mechanisms, with one of the fastest being a shift in and shuffling of their partnerships with symbiotic microorganisms. Assessing symbiosis-focused mechanisms of acclimation and adaptation in response to ocean warming is a technical challenge due to the difficulty of accurately simulating the *de novo* formation of coastal communities. Here, we use the Kiel Outdoor Benthocosm facility to assess which sponges species are experimentally recruited and whether they exhibit symbiosis-focused mechanisms of acclimation following selection to ocean warming. We observed one sponge species (*Haliclona* sp.) and found that this sponge exhibited significant shifts in the membership and composition of its associated microbiome in response to ocean warming, with much of this being attributed to the rare microbiota. Moreover, *Haliclona* sp. maintained the diversity and dominance of its microbiome members. Four bacteria taxa were differentially abundant at elevated temperatures, with two being a *Francisella* sp. that is a suspected pathogen and an uncultured Francisellaceae that is most closely related to sulfur-oxidizing endosymbionts. Changes to the *Haliclona* sp. microbiome are largely consistent with a limited acclimation response, which could indicate that this sponge may use microbial symbionts as part of a mechanism to acclimate and adapt to a warmer future ocean.

## 1. INTRODUCTION

Ocean warming is currently leading to distributional shifts of species towards higher latitudes and colder habitats at a rate of dozens of kilometers per decade (Pinsky et al. 2013, Poloczanska et al. 2013). Due to species-specific differences in thermal window breadths, this will lead to a re-organization of marine communities and cause local changes in biodiversity and associated ecosystem services (García Molinos et al. 2016, Burrows et al. 2019). Anthropogenic perturbations and local change in marine communities are accelerated in bodies of water that are shallow or tidally restricted. The Baltic Sea ecosystem, for example, is a semi-enclosed postglacial sea with a low rate of exchange with the North Atlantic Ocean that exhibits multiple abiotic pressures (*e*.*g*., warming, acidification, and deoxygenation) that mimic those expected for coastal areas in the future (Reusch et al. 2018). With few endemic species due to its young age, it has been proposed that the Baltic Sea can serve as a time-machine to study the ecological and evolutionary impact of anthropogenic perturbations on marine populations (Reusch et al. 2018).

To enable predictions of how distribution ranges may change in the future, it is important to better understand species and community vulnerability to ocean warming, particularly in systems with strong seasonal thermal fluctuations (Wahl et al. 2020). It has been shown that species are often most sensitive to ocean warming during specific life cycle phases or seasons (Pörtner & Farrell 2008, Pandori & Sorte 2019). Coastal communities, for example, are vulnerable to heatwaves during the summer months, with high mortalities observed in a range of systems in the last decades (Reusch et al. 2005, Garrabou et al. 2009). Such mortality could lead to decreased effective population sizes and eventually cause local extinction at the trailing-edge of the thermal window of a given species (Donelson et al. 2019).

Vulnerable species that are most sensitive to ocean warming are able to use several mechanisms to aid in acclimating to environmental change. Much of this capability has been attributed to host physiology. This includes modification of membranes, cellular ultrastructure, and mitochondrial density to compensate for changes in reaction speed of enzymatic reactions (Somero 2022). More recently, acclimation to environmental change has also been attributed to the ability to modify the accessibility of the genome (Eirin-Lopez & Putnam 2019). These means to acclimate and adapt function in concert with an additional mechanism: partnerships with symbiotic microorganisms.

Animals across the tree-of-life have established microbial symbioses that are of deep evolutionary origin and ecological importance (McFall-Ngai et al. 2013). Shifting the membership and composition of microbial partners, regulating microbial abundance, and differential expression of microbial genes all serve as acclimation mechanisms by holobionts (Rosenberg et al. 2009, Carrier & Reitzel 2017). These, in turn, may act as a selective force that enables the holobiont to reach a new healthy state and allow it to better cope with environmental change. This may become a source of adaptation if the microbes are transmitted between generations (Bordenstein & Theis 2015, Webster & Reusch 2017, Voolstra & Ziegler 2020). Shuffling and switching of symbiotic partnerships, in particular, have been shown to increase the thermal tolerance of animals ranging from cnidarians to vertebrates (Baldassarre et al. 2022, Fontaine et al. 2022). Assessing whether microbial symbioses are part of the acclimation response is particularly concerning for animal species that have disproportionate impacts on their communities (*e*.*g*., keystones and ecosystem engineers).

Marine sponges (phylum Porifera) arose >600 million years ago and are among the oldest extant Metazoan lineages (Dunn et al. 2008). The >9,300 species of marine sponges are grouped into four major classes: Calcarea (calcareous sponges; 8%), Demospongiae (demosponges; 83%), Hexactinellida (glass sponges; 7%), and Homoscleromorpha (1%) (Van Soest et al. 2018). These sessile invertebrates filter-feed by pumping thousands of liters of seawater per kilogram of sponge per day. Particulate and dissolved nutrients are then utilized in partnership with symbiotic microorganisms (∼10^6^ to ∼10^9^ cells per mL) that can equate to ∼40% of sponge tissue by volume (Vacelet 1975). This makes sponges vital to benthic-pelagic coupling and the biogeochemical cycling of nutrients in the tropical, temperate, and polar ecosystems that they inhabit (Vogel 1977, Maldonado et al. 2012, Pawlik & McMurray 2020). Marine sponges also form dense aggregations (called sponge gardens or grounds), which can collectively transform the structure of their community by providing distinct habitats that enhance biodiversity (Maldonado et al. 2017).

Assessing symbiosis-focused mechanisms of acclimation and adaptation in response to ocean warming by marine holobionts is a technical challenge due to the difficulty of accurately simulating the *de novo* formation of coastal communities. This is particularly true for sponges, which are notoriously difficult to maintain in laboratory aquaria due to their nutritional requirements and size (Pita et al. 2016). We used the Kiel Outdoor Benthocosms to compensate for these limitations (Figure 1A; Wahl et al. 2015), which has recently been equipped with peristaltic pumps that can transfer planktonic early life stages gently into the experimental units by avoiding the damaging centrifugal acceleration of commonly used rotary pumps. This, in turn, enabled us to recruit from the standing genetic variation of marine species, establish *de novo* communities under ambient and future ocean temperature regimes, and to study how acclimated and adapted genotypes vary between thermal regimes. This colonization included one sponge species (Figure 2A) and, thus, we tested the hypothesis that this sponge has acclimated and adapted to ocean warming by shifting the membership and composition of its microbial partners.

**Figure 1:**
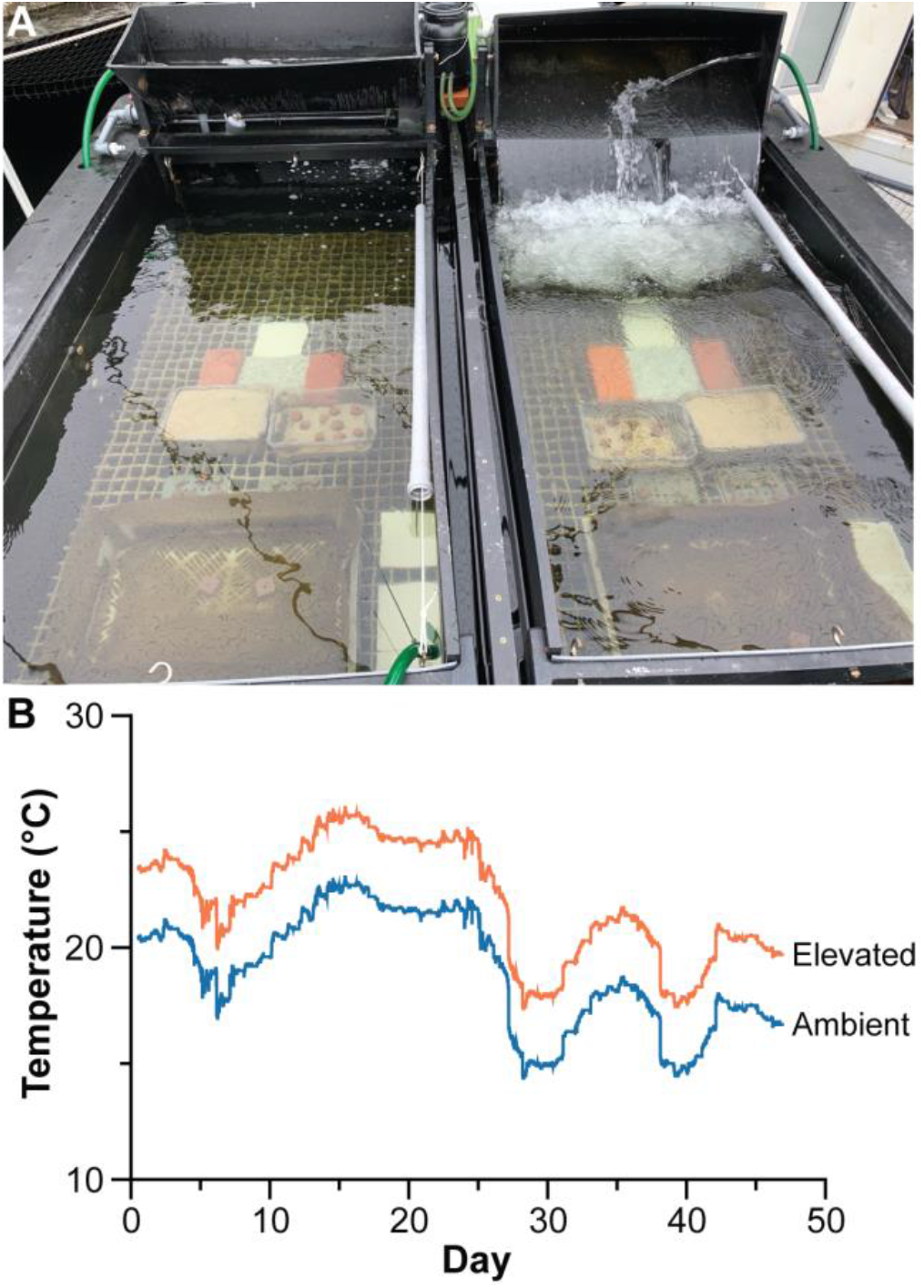
Benthocosms and thermal conditions. (A) Two benthocosm units at the start of the experimental period. Different types of substrates were offered on a GFK-grid (internal openings 3.2 x 3.2 cm) suspended in 30 cm depth (2 small plastic boxes with 1 mm sieved sand and small sand stones, 7 ceramic/stone tiles on the grid and 3 vertically suspended tiles on the right side of each tank, 1 large plastic cage covered with 1 mm plastic mesh and small sand stones, 2 PVC plates with small sand stones attached). The large plastic cage (60 x 40 x 12 cm) was quantitatively surveyed for sponges and tunicates (>5 mm diameter/length) at the end of the experiment. Sponges settled on all surfaces (tank walls, grids and all shaded surfaces of the introduced settlement substrates. (B) Temperature of the seawater in the Kiel Outdoor Benthocosms for ambient (blue) and elevated (orange) tanks.

**Figure 2:**
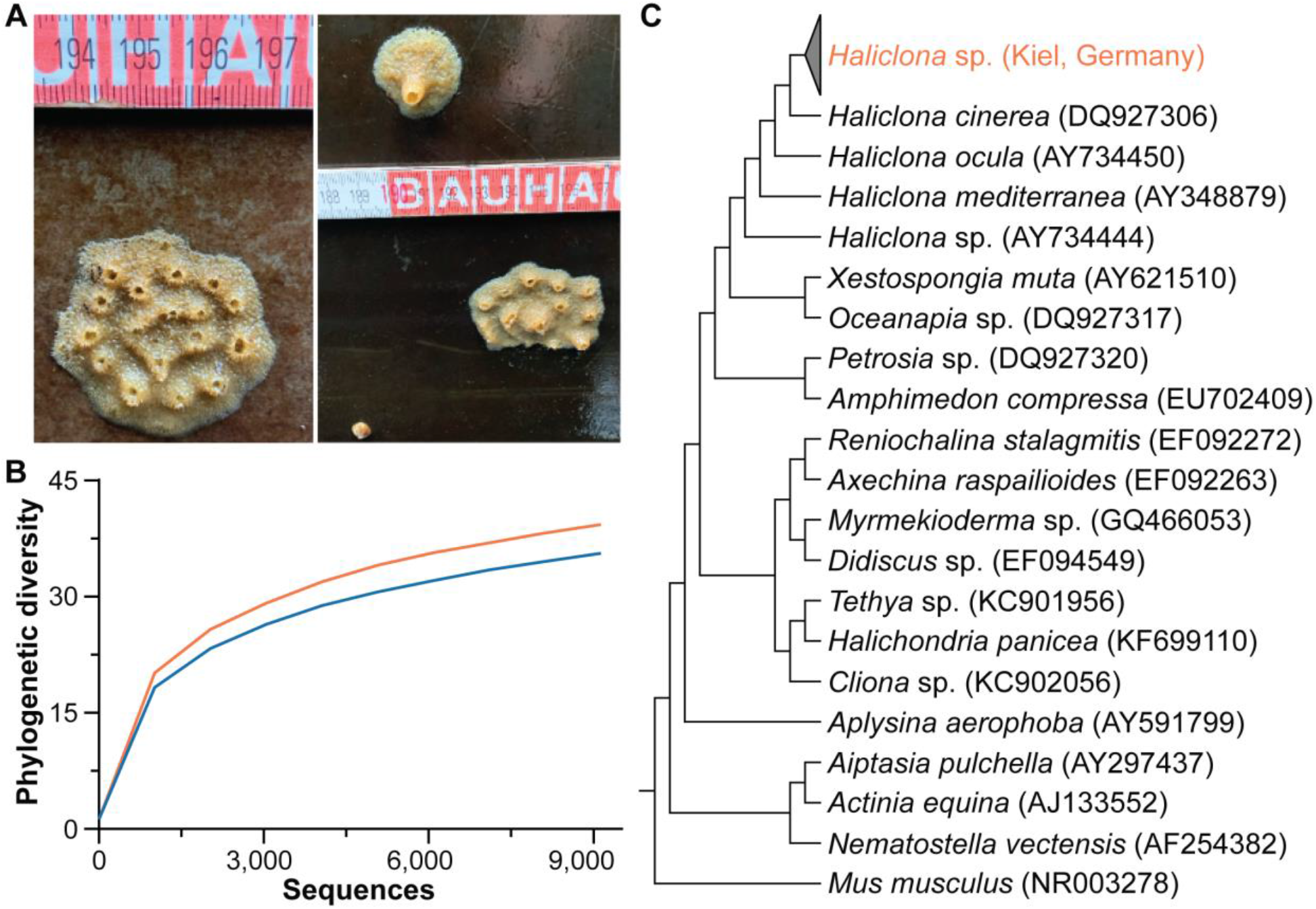
Host identification and microbiome parameters. (A) Sponge (*Haliclona* sp.) colonies that are 3 to 5 cm in diameter on the shaded tank walls of benthocosms during sampling. (B) Rarefaction curves for the phylogenetic diversity of the microbial community associated with *Haliclona* sp. This was based on a rarefaction depth of 9,134 sequences and was used for all analyses. (C) An 18S rRNA phylogeny of *Haliclona* collected from Kiel (Germany) compared with sequences of 19 other demosponges (including 4 *Haliclona* species), mouse *Mus musculus* and sea anemone *Nematostella vectensis* as outgroups.

## 2. MATERIALS AND METHODS

### 2.1. Experimental benthocosms

The Kiel Outdoor Benthocosm facility was used to experimentally determine the colonization potential and microbial ecology of local marine fauna. Briefly, the Kiel Outdoor Benthocosm consists of 12 large (1,500 L) tanks. The tanks are located on a floating dock in the Kiel Fjord (54.330138, 10.149757), where natural, unfiltered seawater can be rapidly pumped from ∼1 m depth into small (5 L) header tanks located on top of each tank using peristaltic pumps (three pumps, each one supplying four experimental tanks that enable transfer of living early life stages from the fjord (Verderflex Dura, Verder GmbH). The experimental tanks are then gravity fed and water is circulated within tanks using pumps and wave generators that create water velocities of 0.2 to 0.4 m/s to ensure rapid mixing of water masses. Surplus water leaves tanks via an overflow.

The water inflow to each of the replicate tanks was set to a rate of approximately 10,000 L/day to supply enough phytoplankton and particulate organic carbon to the benthic communities that colonize each tank. Thus, living larvae can enter the tanks, are selected by environmental and biotic conditions within each tank, settle, and are further selected while acclimating to the respective thermal treatments.

Temperature of each tank was dynamically adjusted using a feedback-control system that uses temperatures continuously measured in the fjord (∼1 m depth) as a baseline (GHL Advanced Technology GmbH, Germany). We randomly assigned one of each of the six replicate benthocosm units as either ambient or elevated (*i.e*., ambient + 3.0 °C to reflect the RCP 8.5-like year 2100 thermal scenarios; Meier et al. 2022). Various amounts of stainless-steel heaters (Schego GmbH, Germany) and chillers (AquaMedic GmbH, Germany) in each of the experimental tanks ensured that the elevated temperature could be maintained throughout the experiment, with maximum offsets between the target and realized temperatures of ∼0.1 °C (Figure 1B). Temperatures were recorded in each of the experimental tanks at 10-minute intervals. The experiment was run between 2 July 2021 and 1 December 2021, with a primary aim to test the effectiveness of the new peristaltic pumps to transfer sensitive larval stages into the Kiel Outdoor Benthocosm tanks (to be published elsewhere).

Tanks were equipped with GFK-grids (mesh size: 32 x 32 mm) that were suspended ∼30 cm below water levels. Multiple structures were placed on these grids in order to provide a diverse set of habitats for competent larvae from various taxa (*i.e*., algae, annalids, cnidarians, crustaceans, echinoderms, mollusks, sponges, and tunicates) to settle on. This ranged from various ceramic and stone tiles to empty and sand filled plastic boxes (Figure 1A). All surfaces of one plastic box (60 x 40 x 12 cm) were analyzed to estimate sponge density in each of the experimental tanks.

### 2.2. Sample collection and species identification

Tissues (*i.e*., 3 technical replications per species per tank) were collected by dissecting ∼2 cm^2^ pieces of tissue from the sponges attached to the surface of each benthocosm. Tissues were gently removed of seawater, preserved in 99% ethanol, and were stored at -20 ºC. Total DNA was extracted from all samples, as well as DNA blanks for contaminant controls (n = 3), according to the manufacturer’s protocol for the DNeasy^®^ Blood & Tissue Mini Kit (Qiagen). Total DNA was quantified using the dsDNA HS Assay Kit for the Qubit Fluorometer (Thermo Fisher Scientific) following the manufacturer’s protocol.

The 18S rRNA gene was amplified according to Redmond et al. (2013) using DreamTaq (Thermo Scientific), isolated using gel electrophoresis, and cleaned using the NucleoSpin Gel and PCR Clean-up kit (Marcherey Nagel). Samples were sent to Eurofins Genomics for Sanger sequencing using their LightRun Service. We performed an initial basic local alignment search tool (BLAST) search on National Center for Biotechnology Information (NCBI) to identify the genus of each sample. We then aligned these sequences to 19 other demosponges (including four others from candidate genus) as well as the mouse *Mus musculus* and sea anemone *Nematostella vectensis* as outgroups using MAFFT to then generate a maximum-likelihood phylogeny using the NGPhylogeny.fr platform (Criscuolo & Gribaldo 2010, Guindon et al. 2010, Katoh & Standley 2013, Lemoine et al. 2019). This phylogenic tree was then visualized using the Interactive Tree Of Life (Letunic & Bork 2021) and stylized using Adobe Illustrator (v. 24.0.1).

### 2.3. Bacterial community analysis

We then diluted the total DNA to 0.5 ng/µL for amplicon sequencing. Samples for amplicon sequencing (V3/V4 region of the 16S rRNA gene) were sent to the Competence Center for Genomic Analysis (Christian-Albrechts University of Kiel, Germany), where libraries were prepared and Illumina MiSeq sequencing (v3, 2×300 bp paired-end reads) was performed.

Raw reads and quality information from the forward sequences were imported into QIIME 2 (v. 2022.11; Bolyen et al. 2019), filtered by quality score, and denoised using Deblur (Amir et al. 2017). QIIME 2-generated ‘features’ were analyzed as amplicon sequence variants (ASVs; Callahan et al. 2017) and were assigned taxonomy using SILVA (v. 138; Quast et al. 2013). Sequences matching to Archaea, chloroplast, mitochondria, and those that were present in the DNA kit blanks were discarded. The filtered table was rarified to 9,134 sequences (Figure 2B). Mean relative abundance for each ASV was then calculated to merge technical replicates.

Unweighted and weighted UniFrac (Lozupone & Knight 2005) distances were calculated in QIIME 2. They were then visualized using Principal Coordinate Analysis (PCoA) in QIIME 2, the *qiime2R* package in R (Bisanz 2018), phyloseq (McMurdie & Holmes 2013), and Adobe Illustrator (v. 24.0.1) for stylization. Permutational analysis of variance (PERMANOVA) and permutational multivariate analysis of dispersion (PERMDISP), and their respective pairwise comparisons, were performed within QIIME 2 to test whether community membership and composition differed between thermal conditions. We then calculated four measures of alpha diversity (total ASVs, Faith’s phylogenetic distance, McIntosh evenness, and McIntosh dominance). Each alpha diversity value was tested for normality using the Shapiro-Wik test and then compared statistically using a two-tailed t-test in Prism (v. 9.0.0). Taxonomy of these communities was then summarized for each thermal condition and the ASVs that were shared and unique to each condition were determined using Venny (v. 2.1.0; Oliveros 2015). Lastly, differentially abundant ASVs were determined using analysis of compositions of microbiomes with bias correction (ANCOM-BC) within QIIME 2 (Lin & Peddada 2020).

### 2.4. Phylogeny of an Francisellaceae ASV

We first re-ran our QIIME 2 pipeline to generate paired-end sequences for the differentially abundant ASVs that were of interest. Sequences were aligned to verify that the single-end differentially abundant ASVs were uniquely identical to the front portion of the paired-end sequence. This was performed to generate sequences that were twice the length and, in turn, provide more accurate phylogenetic information. The paired-end sequence was originally defined as an uncultured Francisellaceae and BLAST further suggested that it is most closely related to the thiotrophic gill endosymbiont of *Bathymodiolus* and other marine bivalves. We compared the sequence of this paired-end sequence ASV to ten of the most related sequences from the BLAST search, as well as diverse animal-associated sulfur-oxidizing symbionts (Tian et al. 2014, Ponnudurai et al. 2017). We then aligned these sequences using MAFFT, cleaned with BMGE, and generated a maximum-likelihood phylogeny using the NGPhylogeny.fr platform (Criscuolo & Gribaldo 2010, Guindon et al. 2010, Katoh & Standley 2013, Lemoine et al. 2019). This phylogenic tree was then visualized using the Interactive Tree Of Life (Letunic & Bork 2021) and stylized using Adobe Illustrator (v. 24.0.1).

## 3. RESULTS

Numerous species colonized the offered habitats within each benthocosm unit (Figure 1A). High densities of mussels (*Mytilus* spp.), clams (*Mya arenaria*), barnacles (*Amphibalanus improvises*), tunicates (*Ciona intestinalis*), and patchy recruitment of sea stars (*Asterias rubens*) were observed, indicating that fragile larval specimen were successfully transported into the experimental tanks alive. Sponge larvae settled on all available shaded hard substrates (from various tiles to plastic surfaces) and reached diameters of >5 cm (Figure 2A). In order to estimate differences in sponge abundance in the two different temperature treatments, we enumerated individuals attached to a large plastic box in each tank. Sponge densities were comparable between both treatments [ambient: 14.7 ± 3.8 standard deviation (range: 12 to 22), elevated: 10.5 ± 3.4 standard deviation (range: 4 to 14)]. Sponge sizes were not analyzed, but large specimen of 3 to 5 cm diameter were found in both treatments (Figure 2A).

Individual sponges collected from benthocosms were most closely related to *Haliclona* sp. (98.1% ± 0.3% sequence similarity). An 18S rRNA gene tree suggests that these sequences form a distinct group that is most closely related to *H. cinerea* and then *H. oculata* (Figure 2C). While these samples were most distinct from a reported *Haliclona* sp. (that was collected from the continental shelf of North America), we have designated these individuals as *Haliclona* sp. because an accurate species designation remains unclear (Figure 2C).

We observed a temperature-induced a shift in the membership and composition of the bacterial community associated with *Haliclona* sp., with individuals from elevated temperatures also having a higher degree of variation than those in ambient conditions (PERMANOVA, unweighted UniFrac: p = 0.001; weighted UniFrac: p = 0. 018; PERMDISP, unweighted UniFrac: p = 0.871; weighted UniFrac: p = 0.001; Figure 3). Temperature, however, did not influence the total number (p = 0.779) or phylogenetic diversity (p = 0.307) of bacteria associated with *Haliclona* sp., as well as the dominance (p = 0.892) or evenness (p = 0.904) of those bacterial taxa (Figure 4).

**Figure 3:**
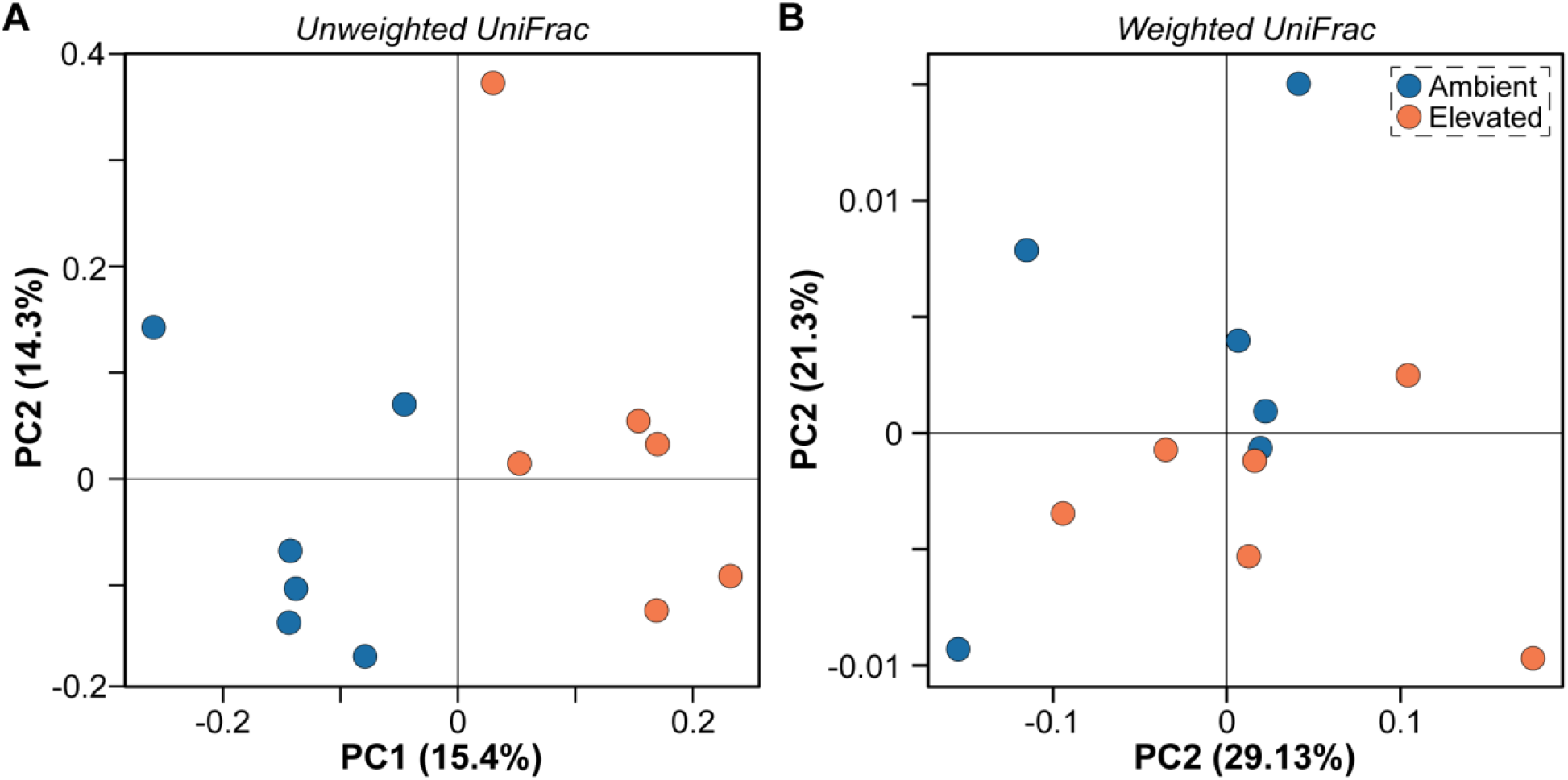
Temperature-induced shift in the microbiome. Temperature induced a shift in the membership (p = 0.001; unweighted UniFrac) and composition (p = 0.018; weighted UniFrac) of the bacterial community associated with *Haliclona* sp.

**Figure 4:**
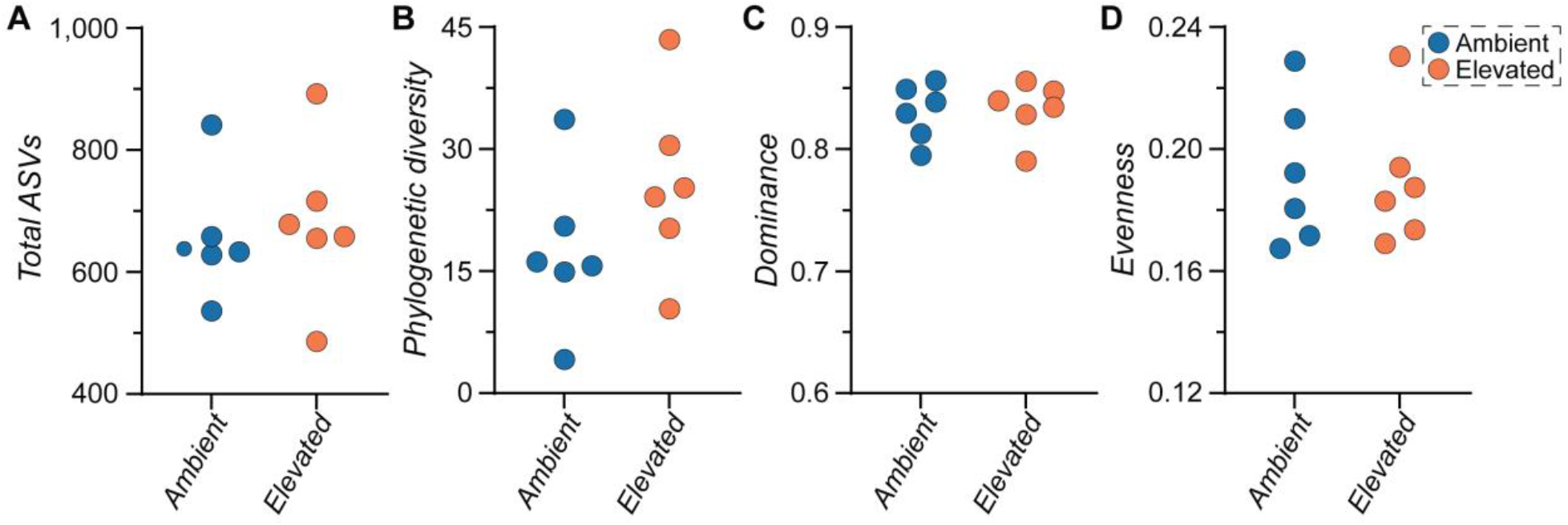
Stability in the *Haliclona* sp. microbiome structure. Temperature did not alter the number of total ASVs (p = 0.779), phylogenetic diversity (p = 0.307), dominance (p = 0.892), or evenness (p = 0.904) (*i.e*., structure) of the bacterial community associated with *Haliclona* sp.

Ten (of the 455) genera represented more than half of the *Haliclona* sp.-associated bacterial community (Figure 5A). The majority (83.6%) of bacterial species representing these genera are shared between both thermal conditions, with 4.9% (representing 0.5% of the of total community) being specific to *Haliclona* sp. in ambient conditions and 11.5% (representing 1.0% of the of total community) being specific to elevated temperatures (Figure 5B). There were 93 bacterial genera that were exclusively observed at elevated temperatures, with 73.2% of these being represented by the *HgCo23* (HgCo23), *NS5* (Flavobacteriales), *PeM15* (PeM15), *Planktomarina* (Rhodobacterales), and *Pseudohongiella* (Oceanospirillales). Those specific to ambient conditions, on the other hand, were mostly (84.2%) represented by *Fulvivirga* (Cytophagales), *SAR86* (SAR86), and an Terasakiellaceae uncultured (Rhodospirillales) (Figure 5B).

**Figure 5:**
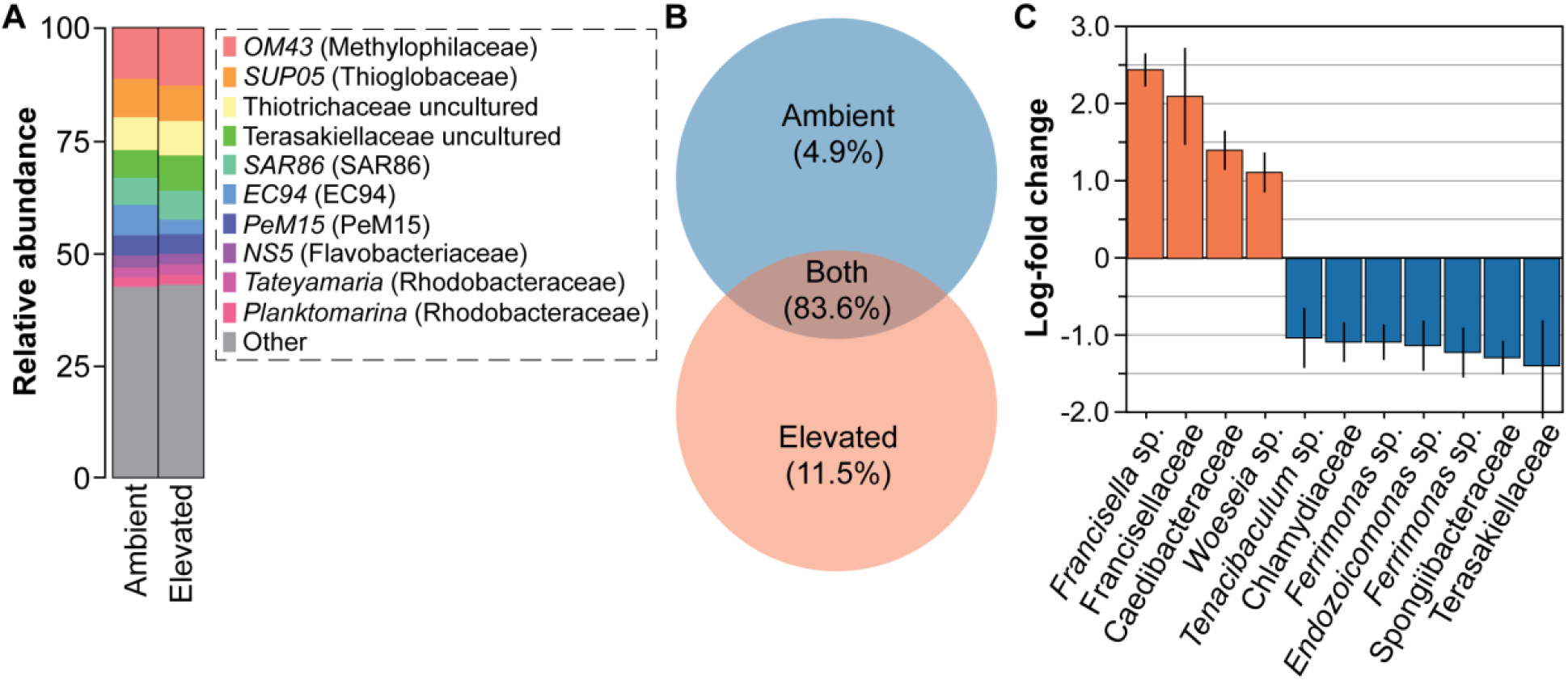
Temperature-induced taxonomic distinctions. (A) Temperature induced slight shifts in the relative abundance for the top 10 most abundant bacterial genera associated with *Haliclona* sp. (B) The majority of bacterial ASVs were shared between thermal conditions for *Haliclona* sp., with ∼12.4% of these being novel associations. (C) Eleven bacterial ASVs were differentially abundant with a log-fold change of more than 1, with four of these being in *Haliclona* sp. from elevated temperatures.

Eleven of the 1,321 total ASVs were differentially abundant between ambient (63.6%) and elevated (36.4%) conditions. ASVs that were differentially abundant in elevated conditions include a *Francisella* sp. (Francisellales), an uncultured Francisellaceae, an uncultured Caedibacteraceae, and a *Woeseia* sp. (Steroidobacterales), which collectively represented 0.6% of the bacterial community in elevated conditions (Figure 5C). Those that were differentially abundant in ambient conditions, on the other hand, include an uncultured Chlamydiaceae (Chlamydiales), Endozoicomonas (Oceanospirillales), two species of *Ferrimonas* (Alteromonadales), an uncultured Spongiibacteraceae (Cellvibrionales), *Tenacibaculum* (Flavobacteriales), an uncultured Terasakiellaceae (Rhodospirillales), which collectively represented 0.8% of the bacterial community in ambient conditions (Figure 5C).

A BLAST search of the uncultured Francisellaceae further suggests that this ASV is within the Thiotrichales. We further determined that the paired-end sequence for this ASV has a ∼99% similarity to the thiotrophic gill endosymbiont of the deep-sea mussel *Bathymodiolus*. A 16S rRNA phylogeny of this *Haliclona* sp. ASV supports that it is most closely related to an uncultured Francisellaceae, Thiotrichales, and bacterioplankton (Figure 6). These four bacteria are then most closely related to the thiotrophic gill endosymbiont of several deep-sea bivalves (*e.g*., *Calyptogena* spp.) and a symbiont of the sponge *Myxilla methanophila* (Figure 6). These are further related to diverse thiotrophic gill endosymbiont of *Bathymodiolus* spp. (Figure 6).

**Figure 6:**
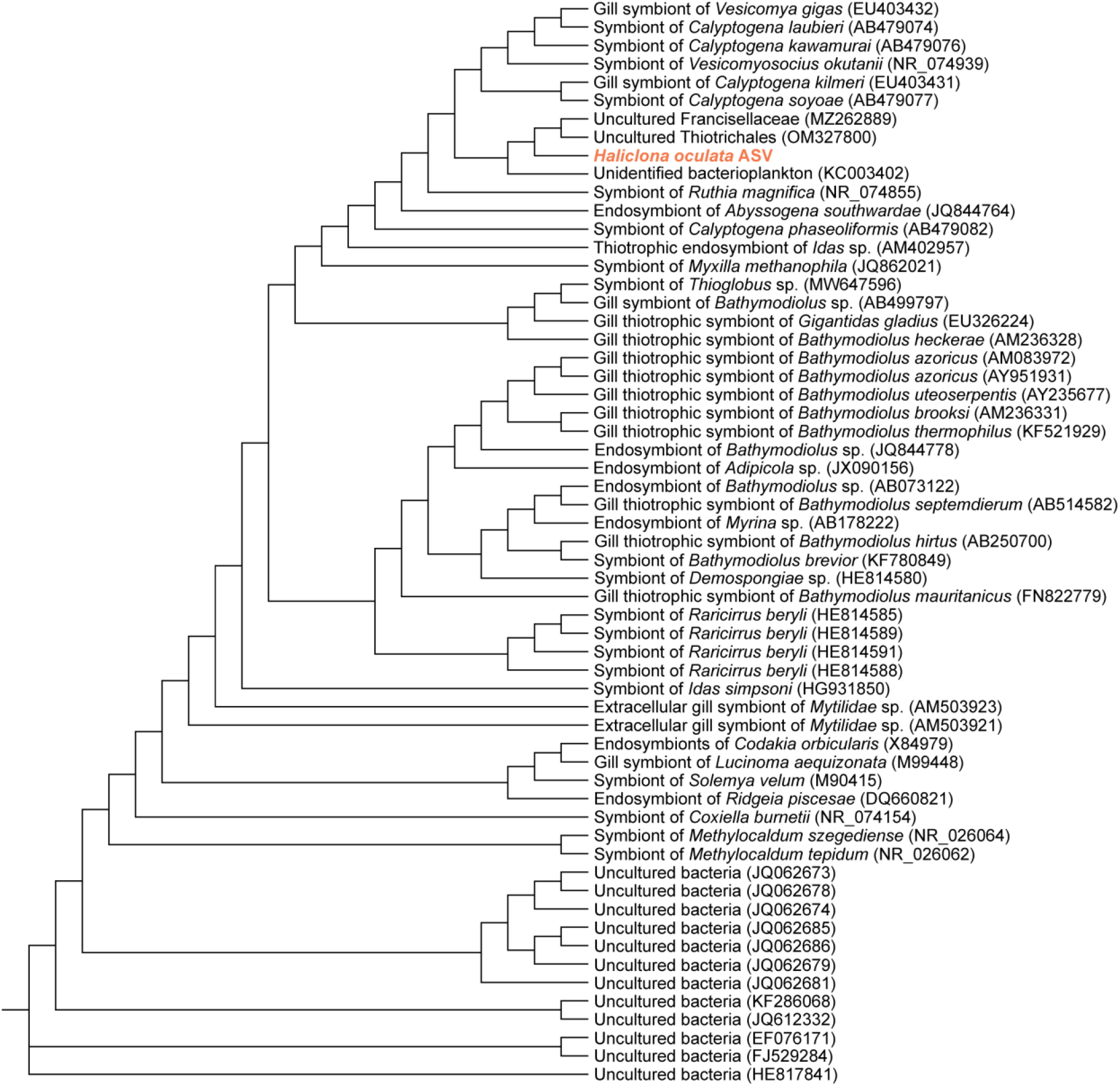
Phylogenetic relationship of uncultured Francisellaceae from *Haliclona* sp. A Maximum Likelihood gene tree for the *Haliclona* sp. ASV-of-interest with a diverse set of sulfur-oxidizing symbionts as well as the ten of the most closely related sequences from BLAST, and distant uncultured bacteria as an outgroup. The *Haliclona* sp. ASV-of-interest is highlighted in red.

## 4. DISCUSSION

Marine sponges associate with hundreds to thousands of microbial taxa, with the composition of these taxa being partially influenced by host evolutionary history and the environment (Thomas et al. 2016, Webster & Thomas 2016, Busch et al. 2022, Pankey et al. 2022). These benthic filter-feeders may shift the membership and composition of their microbial partners or regulate microbial abundance in response to environmental change (Rosenberg et al. 2009, Carrier & Reitzel 2017, Pita et al. 2018). If the environmental change exceeds this acclimation response and adaptive potential, then the sponge-associated microbiome can enter a state of dysbiosis that is often characterized by an increase in the taxonomic and phylogenetic diversity of the symbionts, shift from sponge-enriched microbes to opportunists, and/or a compositional shift of microbial partners in tandem with an increased variation in this compositional profile (Zaneveld et al. 2017, Pita et al. 2018).

Using experimental benthocosms that enabled us to recruit from the standing genetic variation and establish *de novo* coastal communities (Wahl et al. 2015), we observed that the sponge *Haliclona* sp. faced with ocean warming exhibited a significant shift in presence, and to a lesser extent composition, of its associated microbiome, with those shifts being attributed to rare members of this community. Moreover, this sponge did not exhibit a change in the diversity or dominance of the microbiome members, which was similarly observed for the sponge *Carteriospongia foliascens* (Luter et al. 2020). *Haliclona* sp. did, however, show minimal signs of being colonized by opportunists (*i.e*., an *Francisella* that was differentially abundant in elevated temperatures). This response was similar to other sponges, which maintain their association with the most abundant (and, presumably, biologically important) taxa, while there is a shift in the rare microbes (Pita et al. 2018). Thus, when faced with temperatures reflective of ocean warming, data presented here provide minimal support for the premise that changes in the microbiome associated with *Haliclona* sp. is an acclimation response (Rosenberg et al. 2009, Webster & Reusch 2017, Pita et al. 2018).

If *Haliclona* sp. has a limited symbiosis-mediated acclimation potential, then changes in the microbiome may relate to a potential dysbiosis (*e.g*., Lesser et al. 2016). The most differentially abundant bacterial ASV in elevated temperatures was *Francisella*. These bacteria are known for being pathogenic with their role as the causal agent of tularemia being most notable (Keim et al. 2007). This bacterial group, for example, is also a disease and mortality agent of the abalone *Haliotis gigantea* in Japan (Kamaishi et al. 2010, Brevik et al. 2011), the scallop *Patinopecten yessoensis* along the Pacific coast of Canada (Meyer et al. 2017), and the mussel *Mytilus* spp. along the Atlantic coast of France (Garcia et al. 2024). Twenty species of sponges inhabiting coastal communities (and at ambient conditions) have been collectively observed to associate with six species *Francisella* (Thomas et al. 2016). These bacteria collectively represent a small portion of the bacterial community and are most commonly observed in *Carteriospongia foliascens* (Great Barrier Reef, Australia), *Cliona delitrix* (Caribbean Sea), and *Phorbas fictitious* (Portugal) (Thomas et al. 2016). Notably, these *Francisella* species were not detected in *H. tubifera* that was collected from the Caribbean coast of Panama (Thomas et al. 2016). Thus, physiological stress from ocean warming may have led to *Haliclona* sp. to the colonization by and differential abundance of a *Francisella* that may have pathogenic tendencies.

An unexpected and intriguing observation is that the second differentially abundant bacteria that was associated with *Haliclona* sp. in elevated temperatures was an uncultured Francisellaceae that has a nearly identical 16S rRNA sequence to the thiotrophic gill endosymbiont of the deep-sea *Bathymodiolus* and other marine bivalves (Dubilier et al. 2008). These symbionts may be acquired locally in chemically rich environments to then reduced sulphur compounds (Dubilier et al. 2008). Marine sponges act as microbial fermenters to cycle dissolved and particular matter in coastal communities, with sulfur being part of the holobiont metabolism (Hentschel et al. 2006, Hentschel et al. 2012, Webster & Thomas 2016). The sponge *H. cymaeformis* has a sulfur-oxidizing bacterium (*Thioalkalivibrio nitratireducens*) that is localized to the mesohyl to detoxify for the host (Tian et al. 2014). Therefore, in the face of ocean warming, one part of the symbiosis-mediated acclimation response by *Haliclona* sp. may relate to sulfur metabolism within *Haliclona* sp. or the simulated coastal community that it is integrated within.

Our results show signs of ecological changes to the microbiome of *Haliclona* sp. that may also have biological relevance when facing ocean warming. To resolve this, it remains paramount to understand: (i) the functional properties of this microbial community under health and thermal stress, (ii) the genomic potential and expression of this strain of *Francisella* and the uncultured Francisellaceae, and (iii) the physiological response by this species of *Haliclona*. This microbial ecology and holobiont physiology may then be integrated into the context of the ecosystem to help understand, predict, and simulate the direction and fate of benthic communities in the Baltic Sea.

## Data and code availability

Raw amplicon sequencing files have been deposited to the Sequence Read Archive of the NCBI under the BioProject accession number PRJNA1369550. Code corresponding to this amplicon analysis is available on GitHub (https://github.com/TylerJCarrier/HaliclonaMicrobiome_ThermalStress).

## Acknowledgements

We thank Dirk Erpenbeck (Ludwig Maximilian University of Munich, Germany) for phylogenetic confirmation of these *Haliclona* individuals, as well as Erik Borchert and other members of the Research Unit Marine Symbioses at the GEOMAR Helmholtz Centre for Ocean Research (Germany) for comments on an earlier draft of this manuscript.

## Ethical approval

All applicable international, national, and/or institutional guidelines for the care and use of animals were followed. This article does not contain any studies with human participants performed by any of the authors.

## REFERENCES

Amir A, McDonald D, Navas-Molina J, Kopylova E, Morton J, Xu Z, Kightley E, Thompson L, Hyde E, Gonzalez A, Knight R (2017) Deblur rapidly resolves single-nucleotide community sequence patterns. mSystems 2:e00191–00116

Baldassarre L, Ying H, Reitzel A, Franzenburg S, Fraune S (2022) Microbiota mediated plasticity promotes thermal adaptation in the sea anemone *Nematostella vectensis*. Nature Communications 13:3804

Bisanz J (2018) qiime2R: Importing QIIME2 artifacts and associated data into R sessions. https://github.com/jbisanz/qiime2R

Bolyen E, Rideout J, Dillon M, Bokulich N, Abnet C, Al-Ghalith G, Alexander H, Alm E, Arumugam M, Asnicar F, Bai Y, Bisanz J, Bittinger K, Brejnrod A, Brislawn C, Brown C, Callahan B, Caraballo-Rodríguez A, Chase J, Cope E, Da Silva R, Dorrestein P, Douglas G, Durall D, Duvallet C, Edwardson C, Ernst M, Estaki M, Fouquier J, Gauglitz J, Gibson D, Gonzalez A, Gorlick K, Guo J, Hillmann B, Holmes S, Holste H, Huttenhower C, Huttley G, Janssen S, Jarmusch A, Jiang L, Kaehler B, Kang K, Keefe C, Keim P, Kelley S, Knights D, Koester I, Kosciolek T, Kreps J, Langille M, Lee J, Ley R, Liu Y, Loftfield E, Lozupone C, Maher M, Marotz C, Martin B, McDonald D, McIver L, Melnik A, Metcalf J, Morgan S, Morton J, Naimey A, Navas-Molina J, Nothias L, Orchanian S, Pearson T, Peoples S, Petras D, Preuss M, Pruesse E, Rasmussen L, Rivers A, Robeson, II M, Rosenthal P, Segata N, Shaffer M, Shiffer A, Sinha R, Song S, Spear J, Swafford A, Thompson L, Torres P, Trinh P, Tripathi A, Turnbaugh P, Ul-Hasan S, van der Hooft J, Vargas F, Vázquez-Baeza Y, Vogtmann E, von Hippel M, Walters W, Wan Y, Wang M, Warren J, Weber K, Williamson C, Willis A, Xu Z, Zaneveld J, Zhang Y, Zhu Q, Knight R, Caporaso J (2019) Reproducible, interactive, scalable and extensible microbiome data science using QIIME 2. Nature Biotechnol 37:852–857

Bordenstein S, Theis K (2015) Host biology in light of the microbiome: ten principles of holobionts and hologenomes. PLoS Biol 13:e1002226

Brevik Ø, Ottem K, Kamaishi T, Watanabe K, Nylund A (2011) *Francisella halioticida* sp. nov., a pathogen of farmed giant abalone (*Haliotis gigantea*) in Japan. Journal of Applied Microbiology 111:1044–1056

Burrows M, Bates A, Costello M, Edwards M, Edgar G, Fox C, Halpern B, Hiddink J, Pinsky M, Batt R, García Molinos J (2019) Ocean community warming responses explained by thermal affinities and temperature gradients. Nature Clim Change 9:959–963

Busch K, Slaby B, Bach W, Boetius A, Clefsen I, Colaço A, Creemers M, Cristobo J, Federwisch L, Franke A, Gavriilidou A, Hethke A, Kenchington E, Mienis F, Mills S, Riesgo A, Ríos P, Roberts E, Sipkema D, Pita L, Schupp P, Xavier J, Rapp H, Hentschel U (2022) Biodiversity, environmental drivers, and sustainability of the global deep-sea sponge microbiome. Nature Communications 13:5160

Callahan BJ, McMurdie PJ, Holmes SP (2017) Exact sequence variants should replace operational taxonomic units in marker-gene data analysis. The ISME Journal 11:2639–2643

Carrier T, Reitzel A (2017) The hologenome across environments and the implications of a host-associated microbial repertoire. Frontiers in Microbiology 8:802

Criscuolo A, Gribaldo S (2010) BMGE (Block Mapping and Gathering with Entropy): a new software for selection of phylogenetic informative regions from multiple sequence alignments. BMC Evolutionary Biology 10:210

Donelson J, Sunday J, Figueira W, Gaitán-Espitia J, Hobday A, Johnson C, Leis J, Ling S, Marshall D, Pandolfi J, Pecl G (2019) Understanding interactions between plasticity, adaptation and range shifts in response to marine environmental change. Philosophical Transactions of the Royal Society B 374:20180186

Dubilier N, Bergin C, Lott C (2008) Symbiotic diversity in marine animals: the art of harnessing chemosynthesis. Nature Reviews Microbiology 6:725–740

Dunn C, Hejnol A, Matus D, Pang K, Browne W, Smith S, Seaver E, Rouse G, Obst M, Edgecombe G, Sørensen M, Haddock S, Schmidt-Rhaesa A, Okusu A, Kristensen R, Wheeler W, Martindale M, Giribet G (2008) Broad phylogenomic sampling improves resolution of the animal tree of life. Nature 452:745–749

Eirin-Lopez J, Putnam H (2019) Marine environmental epigenetics. Annual Review of Marine Science 11:335–368

Fontaine S, Mineo P, Kohl K (2022) Experimental manipulation of microbiota reduces host thermal tolerance and fitness under heat stress in a vertebrate ectotherm. Nature Ecology & Evolution 6:405–417

Garcia C, Charles M, Chollet B, Nadeau A, Serpin D, Quintric L, Pépin J-F, Houssin M, Lupo C (2024) Understanding the role of *Francisella halioticida* in mussel mortalities in France: an integrative approach. Diseases of Aquatic Organisms 158:81–99

García Molinos J, Halpern B, Schoeman D, Brown C, Kiessling W, Moore P, Pandolfi J, Poloczanska E, Richardson A, Burrows M (2016) Climate velocity and the future global redistribution of marine biodiversity. Nature Climate Change 6:83–88

Garrabou J, Coma R, Bensoussan N, Bally M, Chevaldonné P, Cigliano M, Díaz D, Harmelin J, Gambi M, Kersting D, Ledoux J (2009) Mass mortality in Northwestern Mediterranean rocky benthic communities: effects of the 2003 heat wave. Global Change Biology 15:1090–1103

Guindon S, Dufayard J, Lefort V, Anisimova M, Hordijk W, Gascuel O (2010) New algorithms and methods to estimate maximum-likelihood phylogenies: assessing the performance of PhyML 3.0. Systematic Biology 59:307–321

Hentschel U, Piel J, Degnan S, Taylor M (2012) Genomic insights into the marine sponge microbiome. Nature Reviews Microbiology 10:641–654

Hentschel U, Usher K, Taylor M (2006) Marine sponges as microbial fermenters. FEMS Microbiol Ecol 55:167–177

Kamaishi T, Miwa S, Goto E, Matsuyama T, Oseko N (2010) Mass mortality of giant abalone *Haliotis gigantea* caused by a *Francisella* sp. bacterium. Diseases of Aquatic Organisms 89:145–154

Katoh K, Standley D (2013) MAFFT multiple sequence alignment software version 7: improvements in performance and usability. Mol Biol Evol 30

Keim P, Johansson A, Wagner D (2007) Molecular epidemiology, evolution, and ecology of *Francisella*. Annals of the New York Academy of Sciences 1105:30–66

Lemoine F, Correia D, Lefort V, Doppelt-Azeroual O, Mareuil F, Cohen-Boulakia S, Gascuel O (2019) NGPhylogeny.fr: new generation phylogenetic services for non-specialists. Nucleic Acids Research 47:W260–W265

Lesser M, Fiore C, Slattery M, Zaneveld J (2016) Climate change stressors destabilize the microbiome of the Caribbean barrel sponge, *Xestospongia muta*. Journal of Experimental Marine Biology and Ecology 475:11–18

Letunic I, Bork P (2021) Interactive Tree Of Life (iTOL) v5: an online tool for phylogenetic tree display and annotation. Nucleic Acids Research 49:W293–W296

Lin H, Peddada S (2020) Analysis of compositions of microbiomes with bias correction. Nature Communications 11:3514

Lozupone C, Knight R (2005) UniFrac: a new phylogenetic method for comparing microbial communities. Applied and Environmental Microbiology 71:8228–8235

Luter H, Andersen M, Versteegen E, Laffy P, Uthicke S, Bell J, Webster N (2020) Cross-generational effects of climate change on the microbiome of a photosynthetic sponge. Environmental Microbiology 22:4732–4744

Maldonado M, Aguilar R, Bannister R, Bell J, Conway K, Dayton P, Díaz C, Gutt J, Kelly M, Kenchington E, Leys S (2017) Sponge grounds as key marine habitats: a synthetic review of types, structure, functional roles, and conservation concerns. In: Rossi S, Bramanti L, Gori A, Orejas C (eds) Marine Animal Forests. Springer

Maldonado M, Ribes M, van Duyl F (2012) Nutrient fluxes through sponges: biology, budgets, and ecological implications. Advances in Marine Biology 62:113–182

McFall-Ngai M, Hadfield M, Bosch T, Carey H, Domazet-Loso T, Douglas A, Dubilier N, Eberl G, Fukami T, Gilbert S, Hentschel U, King N, Kjelleberg S, Knoll A, Kremer N, Mazmanian S, Metcalf J, Nealson K, Pierce N, Rawls J, Reid A, Ruby E, Rumpho M, Sanders J, Tautz D, Wernegreen J (2013) Animals in a bacterial world, a new imperative for the life sciences. Proceedings of the National Academy of Sciences 110:3229–3236

McMurdie P, Holmes S (2013) phyloseq: an R package for reproducible interactive analysis and graphics of microbiome census data. PLoS ONE 8:e61217

Meier H, Dieterich C, Gröger M, Dutheil C, Börgel F, Safonova K, Christensen O, Kjellström E (2022) Oceanographic regional climate projections for the Baltic Sea until 2100. Earth System Dynamics Discussions 13:159–199

Meyer G, Lowe G, Gilmore S, Bower S (2017) Disease and mortality among Yesso scallops *Patinopecten yessoensis* putatively caused by infection with *Francisella halioticida*. Diseases of Aquatic Organisms 125:79–84

Oliveros J (2015) Venny. An interactive tool for comparing lists with Venn’s diagrams. https://bioinfogp.cnb.csic.es/tools/venny/index.html

Pandori L, Sorte C (2019) The weakest link: sensitivity to climate extremes across life stages of marine invertebrates. Oikos 128:621–629

Pankey M, Plachetzki D, Macartney K, Gastaldi M, Slattery M, Gochfeld D, Lesser M (2022) Cophylogeny and convergence shape holobiont evolution in sponge–microbe symbioses. Nature Ecology & Evolution 6:750–762

Pawlik J, McMurray S (2020) The emerging ecological and biogeochemical importance of sponges on coral reefs. Annual Review of Marine Science 12:315–337

Pinsky M, Worm B, Fogarty M, Sarmiento J, Levin S (2013) Marine taxa track local climate velocities. Science:1239–1242

Pita L, Fraune S, Hentschel U (2016) Emerging sponge models of animal-microbe symbioses. Frontiers in Microbiology 7:2102

Pita L, Rix L, Slaby B, Franke A, Hentschel U (2018) The sponge holobiont in a changing ocean: from microbes to ecosystems. Microbiome 6:46

Poloczanska E, Brown C, Sydeman W, Kiessling W, Schoeman D, Moore P, Brander K, Bruno J, Buckley L, Burrows M, Duarte C (2013) Global imprint of climate change on marine life. Nat Clim Change 3:919–925

Ponnudurai R, Sayavedra L, Kleiner M, Heiden S, Thürmer A, Felbeck H, Schlüter R, Sievert S, Daniel R, Schweder T, Markert S (2017) Genome sequence of the sulfur-oxidizing *Bathymodiolus thermophilus* gill endosymbiont. Standards in Genomic Sciences 12:50

Pörtner H, Farrell A (2008) Physiology and climate change. Science 322:690–692

Quast C, Pruesse E, Yilmaz P, Gerken J, Schweer T, Yarza P, Peplies J, Glockner F (2013) The SILVA ribosomal RNA gene database project: improved data processing and web-based tools. Nucleic Acids Research 41:590–596

Redmond N, Morrow C, Thacker R, Diaz M, Boury-Esnault N, Cardenas P, Hajdu E, Lôbo-Hajdu G, Picton B, Pomponi S, Kayal E (2013) Phylogeny and systematics of Demospongiae in light of new small-subunit ribosomal DNA (18S) sequences. Integrative and Comparative Biology 53:388–415

Reusch T, Dierking J, Andersson H, Bonsdorff E, Carstensen J, Casini M, Czajkowski M, Hasler B, Hinsby K, Hyytiäinen K, Johannesson K (2018) The Baltic Sea as a time machine for the future coastal ocean. Sci Adv 4:eaar8195

Reusch T, Ehlers A, Hämmerli A, Worm B (2005) Ecosystem recovery after climatic extremes enhanced by genotypic diversity. Proceedings of the National Academy of Sciences 102:2826–2831

Rosenberg E, Sharon G, Zilber-Rosenberg I (2009) The hologenome theory of evolution contains Lamarckian aspects within a Darwinian framework. Environmental Microbiology 11:2959–2962

Somero G (2022) The goldilocks principle: a unifying perspective on biochemical adaptation to abiotic stressors in the sea. Annual Review of Marine Science 14:1–23

Thomas T, Moitinho-Silva L, Lurgi M, Björk J, Easson C, Astudillo-García C, Olson J, Erwin P, López-Legentil S, Luter H, Chaves-Fonnegra A, Costa R, Schupp P, Steindler L, Erpenbeck D, Gilbert J, Knight R, Ackermann G, Lopez J, Taylor M, Thacker R, Montoya J, Hentschel U, Webster N (2016) Diversity, structure and convergent evolution of the global sponge microbiome. Nature Communications 7:11870

Tian R-M, Wang Y, Bougouffa S, Gao Z-M, Cai L, Bajic V, Qian P-Y (2014) Genomic analysis reveals versatile heterotrophic capacity of a potentially symbiotic sulfur-oxidizing bacterium in sponge. Environmental Microbiology 16:3548–3561

Vacelet J (1975) Etude en microscopie electronique de l’association entre bacteries et spongiaires du genre Verongia (Dictyoceratida). Journal de Microscopie et de Biologie Cellulaire 23:271–288

Van Soest R, Boury-Esnault N, Hooper J, Rützler K, De Voogd N, Alvarez de Glasby B, Hajdu E, Pisera A, Manconi R, Schoenberg C, Janussen D (2018) World Porifera Database. The World Register of Marine Species (WoRMS)

Vogel S (1977) Current-induced flow through living sponges in nature. Proceedings of the National Academy of Sciences 74:2069–2071

Voolstra C, Ziegler M (2020) Adapting with microbial help: microbiome flexibility facilitates rapid responses to environmental change. BioEssays 42:2000004

Wahl M, Buchholz B, Winde V, Golomb D, Guy-Haim T, Müller J, Rilov G, Scotti M, Böttcher M (2015) A mesocosm concept for the simulation of near-natural shallow underwater climates: the Kiel Outdoor Benthocosms. Limnology and Oceanography: Methods 13:651–663

Wahl M, Werner F, Buchholz B, Raddatz S, Graiff A, Matthiessen B, Karsten U, Hiebenthal C, Hamer J, Ito M, Gülzow E (2020) Season affects strength and direction of the interactive impacts of ocean warming and biotic stress in a coastal seaweed ecosystem. Limnology and Oceanography 65:807–827

Webster N, Reusch T (2017) Microbial contributions to the persistence of coral reefs. The ISME Journal 11:2167–2174

Webster N, Thomas T (2016) The sponge hologenome. mBio 7:e00135–00116

Zaneveld J, McMinds R, Vega Thurber R (2017) Stress and stability: applying the Anna Karenina principle to animal microbiomes. Nature Microbiology 2:17121

